# Development of a QSAR model to predict molecular inhibition of human STAT3

**DOI:** 10.1101/2021.10.29.466511

**Authors:** Joshua M. Patterson, Kimberly Milligan, Cherese Winstead

## Abstract

Herein is described a method to develop a QSAR model to predict the molecular inhibition of human STAT3. The model was built using machine learning using Python 3.3. A random forest machine learning algorithm was deployed. The model produced a R^2^ value of 0.86, indicating that our QSAR model is effective at predicting inhibitory compounds of STAT3.

## INTRODUCTION

IBD is a set of idiopathic chronic relapsing immune-mediated inflammatory diseases defined by two primary types of intestinal inflammation: ulcerative colitis (UC) and Crohn’s disease (CD) [(1)]. Although the specific processes of IBD pathogenesis are still unknown, research has indicated that several cytokines, including interleukins (ILs) and interferons, are implicated in the disease’s etiology. [(2),(3)] The majority of these cytokines activate members of a family of cytoplasmic transcription factors such as signal transducers and activators of transcription (STATs) [(4)]. Numerous recent investigations on the involvement of STATs in IBD have focused on this topic. Recent research from human IBD studies [(5)] and experimental IBD models [(6)] have shown that STAT3 plays an important role in IBD. These investigations have demonstrated that STAT3 has a unique function in the etiology of IBD because it has different roles in innate and acquired immune cells.

STAT3 was discovered to be an acute phase response factor, an inducible DNA binding protein that binds to the IL-6 responsive element inside the promoters of hepatic acute phase protein genes. [(7)] STAT3 is activated in response to a wide range of cytokines and growth factors, according to subsequent research. [(8)] STAT3 is immediately phosphorylated after these cytokines and growth factors ligate their specific receptors, resulting in dimerization and migration to the nucleus to induce several gene expressions encoding molecules that play a role in a variety of biological functions such as cell growth, anti- and pro-apoptosis, cytokine proliferation activation, and cell motility. Besides IBD, STAT3 has also been implicated in playing a major role in the sustaination of certain cancers. This makes STAT3 an ideal target for inhibition. The purpose of this study, therefore, is to identify potential small-molecule inhibitors of STAT3 via QSAR modeling using machine learning.

## MATERIALS AND METHODS

A summary of the workflow of this study is provided in Figure 1. Briefly, a large-scale QSAR model for predicting and analyzing STAT3 inhibition was developed using machine learning; the model was built using a Jupyter notebook powered by Python 3.6; the Jupyter notebook add-in in Visual Studio Code (version number) was used. The code is based upon the work by Chanin Nantasenamat [(9)].

**Figure 1:**
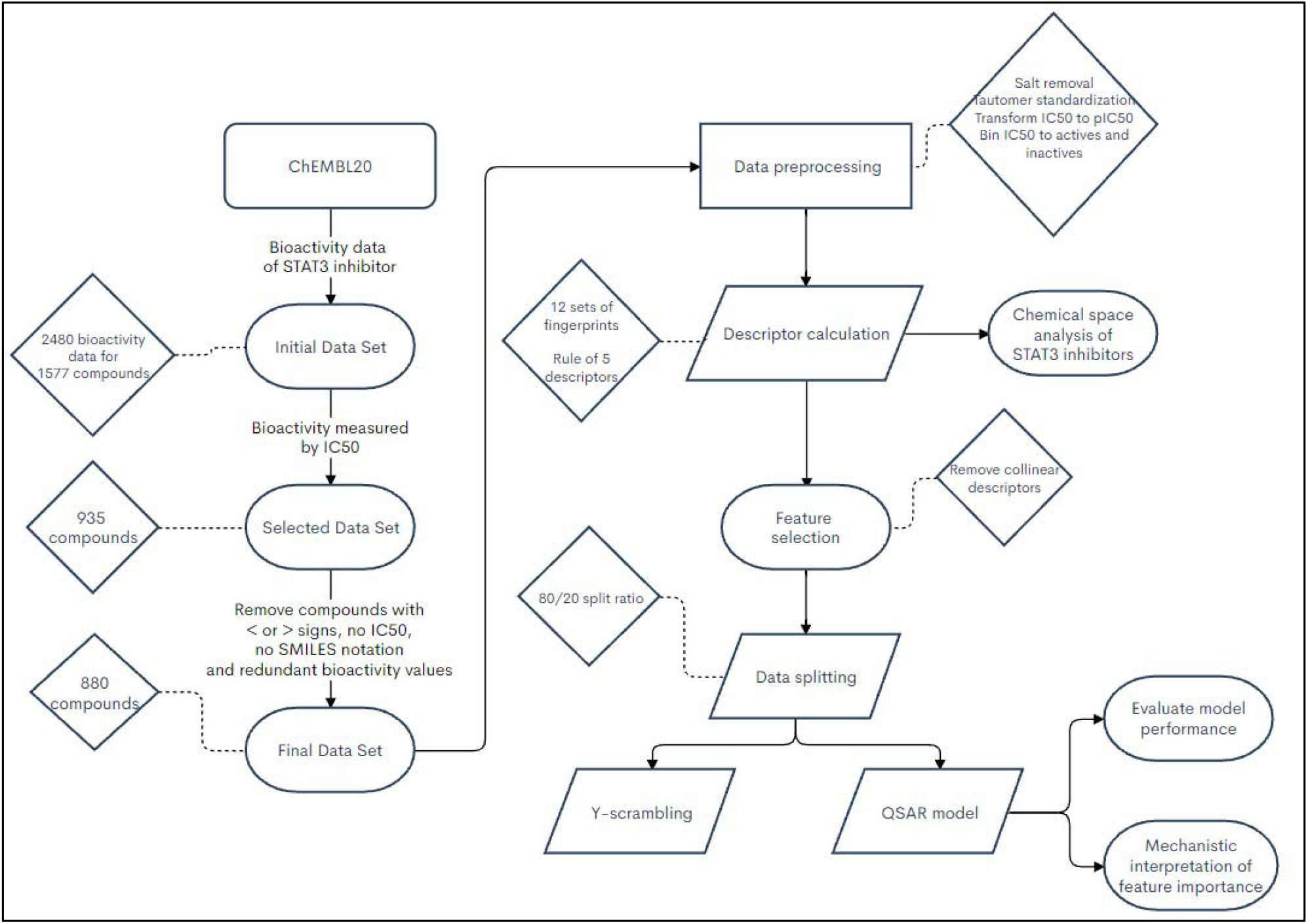
Workflow of QSAR modeling and molecular docking for investigating STAT3 inhibitory activity.

### Data set

A data set of inhibitors against human STAT3 (Target ID CHEMBL4026), comprised of a total number of 2,480 bioactivity data points from 1,577 compounds, were compiled from the ChEMBL20 database; the data was retrieved using the ChEMBL webresource client database add-in pip installed into a Jupyter notebook.

The initial data set was assembled from several bioactivity measurement units including IC_50_, *K*_*i*_, % activity, etc. IC_50_ – the half maximal inhibitory concentration – was selected for further investigation as they constituted the largest subset with 935 compounds. Compounds whose bioactivity data was redundant, or had greater than/less than signs, no IC_50_ value, or no SMILES notation were removed, leaving a final data set comprising of 880 compounds.

### Description of inhibitors

STAT3 inhibitors were encoded by a vector of fingerprint descriptors accounting for its molecular constituents. The 12 molecular fingerprints calculated by the PaDEL-Descriptor software were calculated using Python code [(9)]. The code was written to remove salts and standardize tautomers prior to descriptor calculation. Table 1 summarizes the employed fingerprints along with their corresponding size, description and reference.

**Table 1:**
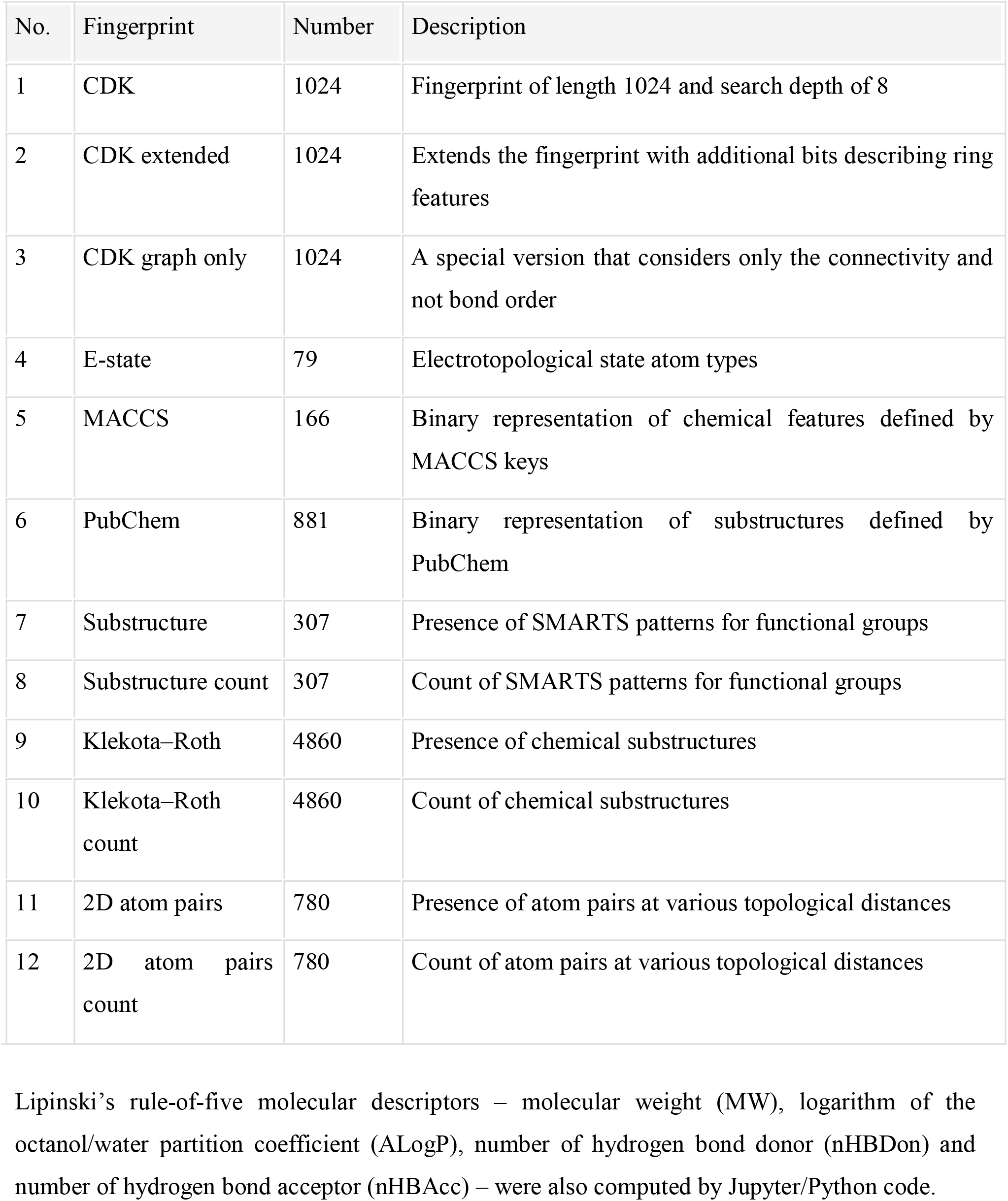
Summary of 12 sets of fingerprint descriptors employed in this study.

### Data splitting

To minimize the possibility of bias that may arise, predictive models were constructed from 100 independent data splits. The data set was split 80/20 into internal and external sets.

### Multivariate analysis

The purpose of supervised learning is to create a model using labeled training data that can be used to predict unknown or future data. This study creates regression models that allow for the prediction of a continuous response variable (pIC50) as a function of predictors (i.e., fingerprint descriptors).

A random forest (RF) is a classifier ensemble composed of numerous decision trees. In a nutshell, the main idea underlying RF is that instead of constructing a deep decision tree with an ever-increasing number of nodes, which might lead to data overfitting and overtraining, multiple trees are built to reduce variance rather than enhance accuracy. As a result, the outputs will be noisier than those of a well-trained decision tree, but they are typically trustworthy and durable.

### Validation of QSAR models

Model validation is a critical step that should be taken to guarantee that a fitted model can effectively predict responses for future or unknown subjects. The performance of the QSAR models was evaluated using two statistical parameters: Pearson’s correlation coefficient (r) and root mean squared error (RMSE). The r value is a popular statistic for expressing the degree of association between two variables of interest. It can vary from 1 to +1, with negative values indicating a negative correlation between two variables and positive values indicating a positive correlation between two variables. The root mean square error (RMSE) is a regularly used measure to analyze the prediction model’s relative error.

The QSAR models’ predictive ability was validated using 10-fold cross-validation, external validation, and the Y-scrambling test. To develop a predictive model, the 10-fold cross-validation approach does not employ the complete data set. Instead, it divides the data into training and testing data sets, allowing the model developed using the training data set to be evaluated on the testing data set. The average accuracies may be utilized to genuinely analyze the prediction model’s performance by repeating the 10-fold validation.

## RESULTS AND DISCUSSION

### Chemical space of STAT3 inhibitors

The Lipinski’s rule-of-five descriptors were used to navigate the chemical space of STAT3 inhibitors in order to get insights into the structure–activity connection. Chemical space analysis can give valuable information about the general features of compounds that determine their inhibitory activities. Lipinski’s rule-of-five descriptors (MW, ALogP, nHBDon, and nHBAcc) were used for exploratory data analysis. The molecular weight (MW) of a chemical is often employed because it is easy to read and compute, and the right size of a drug is critical for its transit across a lipid membrane.

ALogP is a frequently used measure for assessing a compound’s lipophilicity and for estimating membrane penetration and permeability. nHBDon and nHBAcc denote the number of hydrogen bond donors and acceptors, which are used to calculate hydrogen bonding capacity.

The chemical space of ALogP as a function of MW is shown in Figure 2, as to investigate the chemical space of STAT3 inhibitors. A dense distribution of inhibitors was observed within the space of MW starting from approximately 200–600 Da and within the space of ALogP ranging from approximately 2.0 to 6.5 for inactives and 500–700 Da and within the space of ALogP ranging from approximately -4.0 to 1.5 for actives.

**Figure 2:**
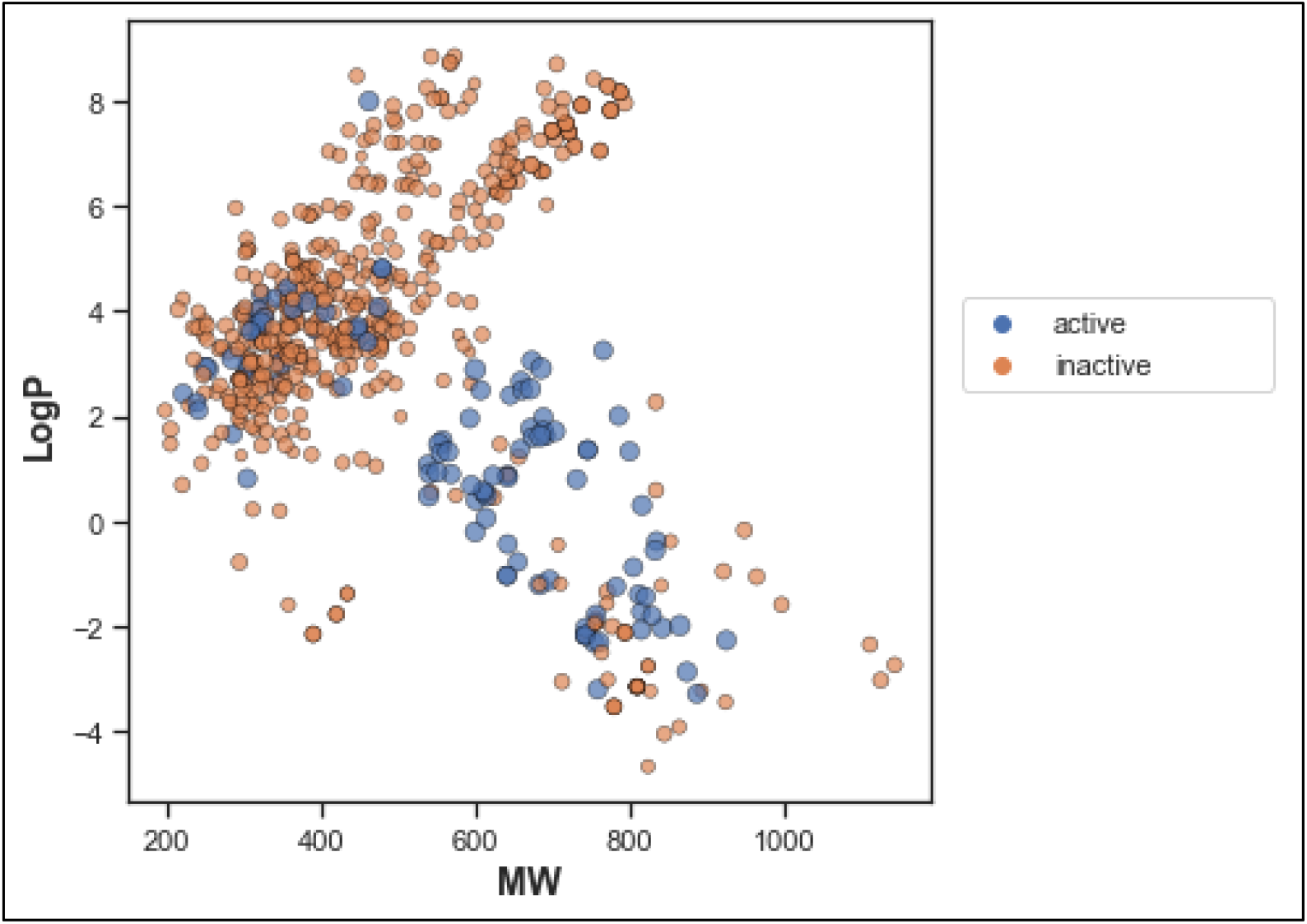
Chemical space of STAT3 inhibitors. Actives and inactives are shown in blue and orange colors, respectively.

The box plot of the Lipinski’s descriptors is shown in Figure 3. Compounds with negative ALogP values approximately of higher vales (above 2) can be found in inactive inhibitors whereas most of the active inhibitors tend to possess approximately lower values in average of ALogP values.

**Figure 3:**
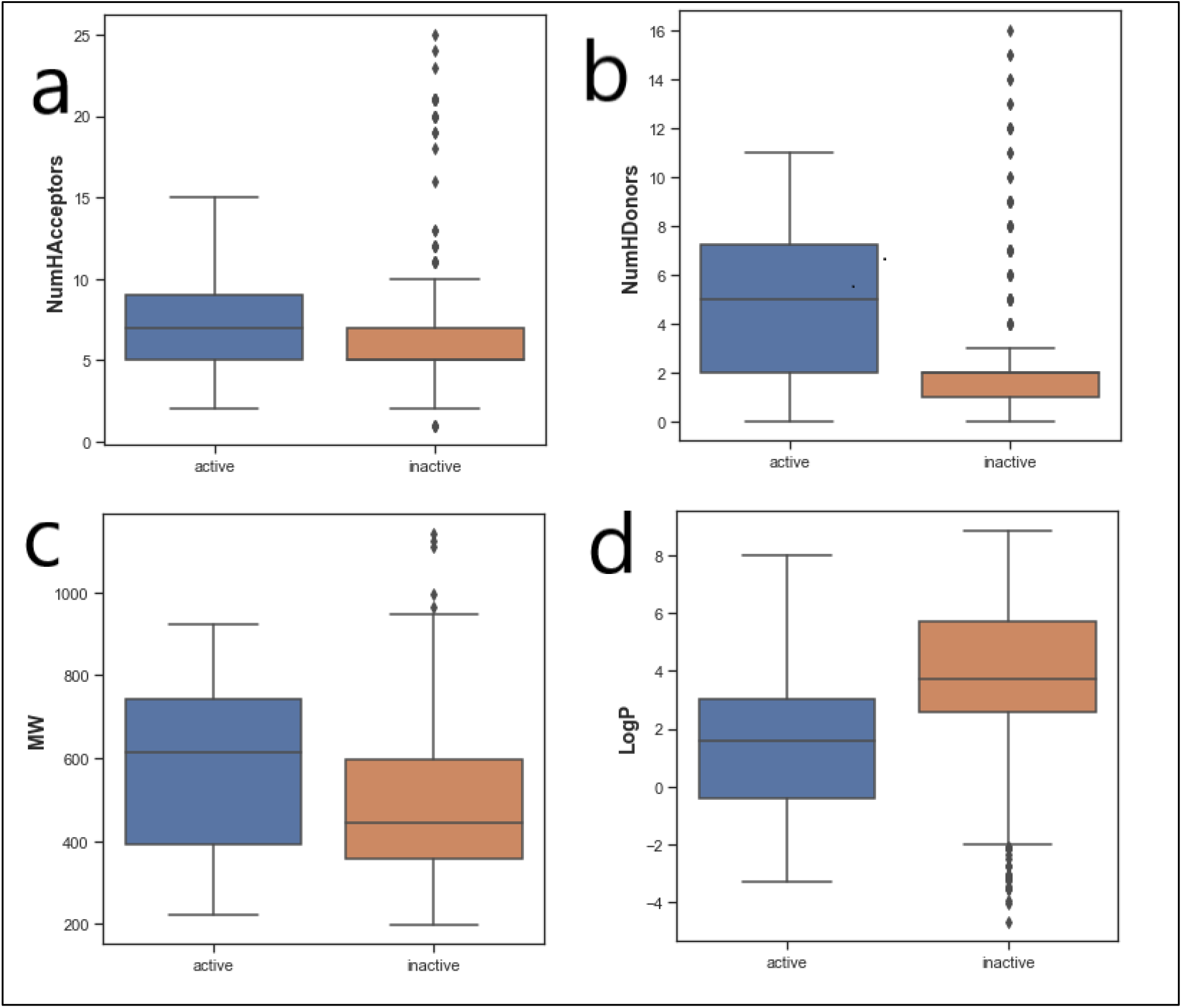
Box plot of STAT3 inhibitors using Lipinski’s rule-of-five descriptors.

### QSAR model for predicting STAT3 inhibitory activity

For the development of QSAR models, a data collection of 880 chemicals was employed. Twelve sets of fingerprint descriptors were specifically benchmarked in order to discover the top performing set. Prior to modeling, collinear descriptors were removed using feature selection. Each of the twelve models was then generated using an 80/20 data split ratio, with 80 percent of the data set serving as the internal set and 20 percent serving as the exterior set.

All twelve models can capture the inhibitory activity space of STAT3 inhibitors since they gave R^2^ values greater than 0.6, indicating robust model performance. The best model was the random forest model built from the Substructure descriptor, which yielded a R^2^ value of 0.86.

Feature importance analysis help reveal features that are important toward bioactivity. Table 2 lists the substructure fingerprints along with their respective descriptions.

**Table 2:** List of top substructure fingerprints and their corresponding description.

| FP | Description |
| --- | --- |
| SubFP1 | Primary carbon |
| SubFP2 | Secondary carbon |
| SubFP3 | Tertiary carbon |
| SubFP5 | Alkene |
| SubFP12 | Alcohol |
| SubFP14 | Secondary alcohol |
| SubFP18 | Alkylarylether |
| SubFP88 | Carboxylic acid derivative |
| SubFP99 | Primary amide |
| SubFP100 | Secondary amide |
| SubFP101 | Tertiary amide |
| SubFP127 | Peptide middle |
| SubFP137 | Vinylogous amide |
| SubFP143 | Carbonic acid derivatives |
| SubFP181 | Hetero N nonbasic |
| SubFP184 | Heteroaromatic |
| SubFP246 | Phosphoric acid derivative |
| SubFP274 | Heterocyclic |
| SubFP275 | Epoxide |
| SubFP287 | Conjugated triple bond |
| SubFP295 | Charged |
| SubFP300 | 1,5-Tautomerizable |
| SubFP301 | Rotatable bond |
| SubFP302 | Michael acceptor |
| SubFP303 | Dicarbodiazene |
| SubFP307 | Chiral center unspecified |

## CONCLUSION

Twelve sets of fingerprint descriptors were utilized to build QSAR models, and their performance was compared. Several fingerprint descriptors had good performance for the built models, indicating that they might capture the feature space of STAT3 inhibitors.

## REFERENCES

1. Podolsky DK. Inflammatory Bowel Disease. N Engl J Med. 1991 Sep 26;325(13):928–37.

2. Bouma G, Strober W. The immunological and genetic basis of inflammatory bowel disease. Nat Rev Immunol. 2003 Jul;3(7):521–33.

3. Mudter J, Neurath MF. Il-6 signaling in inflammatory bowel disease: Pathophysiological role and clinical relevance: Inflamm Bowel Dis. 2007 Aug;13(8):1016–23.

4. Rawlings JS, Rosler KM, Harrison DA. The JAK/STAT signaling pathway. J Cell Sci. 2004 Mar 15;117(8):1281–3.

5. Lovato P, Brender C, Agnholt J, Kelsen J, Kaltoft K, Svejgaard A, et al. Constitutive STAT3 Activation in Intestinal T Cells from Patients with Crohn’s Disease. J Biol Chem. 2003 May;278(19):16777–81.

6. Atreya R, Mudter J, Finotto S, Müllberg J, Jostock T, Wirtz S, et al. Blockade of interleukin 6 trans signaling suppresses T-cell resistance against apoptosis in chronic intestinal inflammation: Evidence in Crohn disease and experimental colitis in vivo. Nat Med. 2000 May;6(5):583–8.

7. Acute-phase response factor, a nuclear factor binding to acute-phase response elements, is rapidly activated by interleukin-6 at the posttranslational level [Internet]. [cited 2021 Oct 29]. Available from: https://journals.asm.org/doi/epdf/10.1128/mcb.13.1.276-288.1993

8. Sugimoto K. Role of STAT3 in inflammatory bowel disease. World J Gastroenterol WJG. 2008 Sep 7;14(33):5110–4.

9. Nantasenamat C. code [Internet]. 2021 [cited 2021 Oct 29]. Available from: https://github.com/dataprofessor/code/blob/027e657f961692e685aff038f73122e68dc9ad1c/python/CDD_ML_Part_1_Acetylcholinesterase_Bioactivity_Data_Concised.ipynb

